# Automated skull stripping in mouse fMRI analysis using 3D U-Net

**DOI:** 10.1101/2021.10.08.462356

**Authors:** Guohui Ruan, Jiaming Liu, Ziqi An, Kaiibin Wu, Chuanjun Tong, Qiang Liu, Ping Liang, Zhifeng Liang, Wufan Chen, Xinyuan Zhang, Yanqiu Feng

## Abstract

Skull stripping is an initial and critical step in the pipeline of mouse fMRI analysis. Manual labeling of the brain usually suffers from intra- and inter-rater variability and is highly time-consuming. Hence, an automatic and efficient skull-stripping method is in high demand for mouse fMRI studies. In this study, we investigated a 3D U-Net based method for automatic brain extraction in mouse fMRI studies. Two U-Net models were separately trained on T2-weighted anatomical images and T2*-weighted functional images. The trained models were tested on both interior and exterior datasets. The 3D U-Net models yielded a higher accuracy in brain extraction from both T2-weighted images (Dice > 0.984, Jaccard index > 0.968 and Hausdorff distance < 7.7) and T2*-weighted images (Dice > 0.964, Jaccard index > 0.931 and Hausdorff distance < 3.3), compared with the two widely used mouse skull-stripping methods (RATS and SHERM). The resting-state fMRI results using automatic segmentation with the 3D U-Net models are identical to those obtained by manual segmentation for both the seed-based and group independent component analysis. These results demonstrate that the 3D U-Net based method can replace manual brain extraction in mouse fMRI analysis.

## 1 Introduction

Functional magnetic resonance imaging (fMRI) (D’Esposito et al., 1998; Lee et al., 2013) has been widely employed in neuroscience research. The key advantage of mouse fMRI (Jonckers et al., 2011; Mechling et al., 2014; Perez-Cervera et al., 2018; Wehrl et al., 2014) is that it can be combined with neuromodulation techniques (e.g., optogenetics) and allows manipulation and visualization of whole-brain neural activity in health and disease, which builds an important link between pre-clinical and clinical research (Lake et al., 2020; Lee et al., 2010; Zerbi et al., 2019). In mouse fMRI research, structural and functional images are commonly acquired with T2-weighted (T2w) and T2*-weighted (T2*w) scanning, respectively. Generally, functional T2*w images need to be registered to standard space using two spatial transformations, which are obtained by registering functional images to anatomical images and subsequently registering anatomical images to a standard space. To exclude the influence of non-brain tissues on image registration, it is necessary to perform skull stripping on both structural and functional images. In the practice of mouse fMRI analysis, skull stripping is usually performed by manually labeling each MRI volume slice-by-slice, due to the absence of a reliable automatic segmentation method. This manual brain extraction is extremely time-consuming, as a large number of slices need to be processed in the fMRI analysis for each mouse. In addition, manual segmentation suffers from intra- and inter-rater variability. Therefore, a fully automatic, rapid, and robust skull-stripping method for both T2w and T2*w images is highly desirable in mouse fMRI studies.

In human research, a number of automatic brain-extraction methods have been developed and widely used, including Brain Extraction Tool (BET) (Smith, 2002), Hybrid Watershed Algorithm (HWA) (Segonne et al., 2004), the Brain Extraction based on nonlocal Segmentation Technique (BEaST) (Eskildsen et al., 2012), and the Locally Linear Representation-based Classification (LLRC) for brain extraction (Huang et al., 2014). However, these methods cannot directly be applied for mouse skull stripping. Compared to human brain MR images, mouse counterparts have relatively lower tissue contrast and a narrower space between the brain and skull, which substantially increases the difficulty of mouse brain segmentation. In addition, the T2*w images used for mouse fMRI may suffer from severe distortion and low spatial resolution, making the skull stripping of functional images more challenging than that for structural images.

Several methods have been proposed for rodent brain extraction. 3D Pulse-Coupled Neural Network (PCNN)-based skull stripping (Chou et al., 2011) is an unsupervised artificial 3D network approach that iteratively groups adjacent pixels with similar intensity and performs morphological operation to obtain the rodent brain mask. Rodent Brain Extraction Tool (T. Wood, 2013) is adapted from the well-known BET (Smith, 2002) method with an appropriate shape for the rodent. Rapid Automatic Tissue Segmentation (RATS) (Oguz et al., 2014) consists of two stages: grayscale mathematical morphology and LOGISMOS-based graph segmentation (Yin et al., 2010), and it incorporates the prior of rodent brain anatomy in the first stage. SHape descriptor selected Extremal Regions after Morphologically filtering (SHERM) (Liu et al., 2020) is an atlas-based method that relies on the fact that the shape of the rodent brain is highly consistent across individuals. This method adopts morphological operations to extract a set of brain mask candidates that match the shape of the brain template, and then merges them for final skull stripping. One of the common disadvantages of the above methods is that their effectiveness was verified only on anatomical images, and cannot be guaranteed on functional images. Another limitation is that the performance of these methods is severely affected by the brain shape, texture, signal to noise ratio, and contrast of images, and hence they need to be optimized for different MRI images. Therefore, it is necessary to develop an automatic skull-stripping method that is effective and has a stable performance on varying types of MR images.

Deep learning has gained popularity in varying image analysis tasks, such as organ or lesion segmentation (Guo et al., 2019; Li et al., 2020; Sun et al., 2019), and disease diagnosis (De Fauw et al., 2018; Suk et al., 2017), owing to its excellent performance. As for skull stripping, Kleesiek et al. (2016) first proposed a 3D convolutional neural network (CNN) method for human brain MR images. Roy et al. (2018) trained a CNN architecture with modified Google Inception (Szegedy et al., 2015) using multiple atlases for both human and rodent brain MR images. With the development of deep learning, more advanced architectures of CNN for semantic segmentation have been proposed. As a popular architecture of deep CNN, U-Net has proved to be very effective in the task of semantic segmentation even with a limited amount of annotated data (Ronneberger et al., 2015). Some U-Net based methods for rodent skull stripping have been proposed. Thai et al. (2019) utilized 2D U-Net for mouse skull stripping on diffusion weighted images. Hsu et al. (2020) trained a 2D U-Net based model using both anatomical and functional brain images for mouse and rat skull stripping. De Feo et al. (2021) proposed a multi-task U-Net to accomplish both skull stripping and brain region segmentation simultaneously on mouse anatomical images of the mouse brain. These methods were only evaluated using segmentation accuracy with reference to the ground truth mask. However, the effect of skull-stripping algorithm on the final fMRI results remains unexplored, and whether automatic skull stripping can replace the manual approach in mouse fMRI analysis remains an open question.

Here, we investigated the feasibility of using 3D U-Net to extract the mouse brain from T2w anatomical and T2* functional images for the fMRI analysis. We separately trained the U-Net models on anatomical T2w and functional T2*w images, considering that anatomical and functional images are acquired with different sequences and have different contrast, resolution, and artifacts. The performance of trained 3D U-Net models is first quantitatively evaluated using conventional accuracy metrics. In addition, we compared the fMRI results separately obtained using the manual and automatic segmentation masks.

## 2 Materials and Methods

### 2.1 Datasets

This study includes two different in-house datasets. Both of these two datasets were collected in the fMRI study and were reanalyzed for the purpose of the present study. All animal experiments were approved by local Institutional Animal Care and Use Committee.

The first dataset (D1) was acquired from 84 adult male C57BL/6 mice (25–30 g) on a Bruker 7.0T MRI scanner using cryogenic RF surface coils (Bruker, Germany). Both the anatomical data (T2w) and resting-state fMRI data (T2*w) were acquired for each mouse. The T2w images were acquired using a fast spin echo (TurboRARE) sequence: field of view (FOV) = 16 × 16 mm^2^, matrix = 256 × 256, in-plane resolution = 0.0625 × 0.0625 mm^2^, slice number = 16, slice thickness = 0.5 mm, RARE factor = 8, TR/TE = 2500 ms/35 ms, number of averages = 2. The resting-state fMRI images were then acquired using a single-shot gradient-echo-planar-imaging sequence with 360 repetitions: FOV = 16 × 16 mm^2^, matrix size = 64 × 64, in-plane resolution= 0.25 × 0.25 mm^2^, slice number = 16, slice thickness = 0.4 mm, flip angle = 54.7°, TR/TE = 750 ms/15 ms.

The second dataset (D2) was obtained from the previous task-state fMRI research (Chen et al., 2020). A total of 27 adult male C57BL/6 mice (18–30 g) were used in this study (Part 1: 13 for auditory stimulation; Part 2: 14 for somatosensory stimulation). The anatomical T2w images and two sets of functional T2*w images were acquired for each mouse with the Bruker 9.4T scanner. The T2w images were acquired using a TurboRARE sequence: FOV = 16 × 16 mm^2^, matrix = 256 × 256, in-plane resolution = 0.0625 × 0.0625 mm^2^, slice number = 32, slice thickness = 0.4 mm, RAREfactor = 8, TR/TE = 3200 ms/33 ms. Two sets of functional T2*w data, acquired using a single-shot echo planar imaging (EPI) sequence, consist of a high spatial resolution one (EPI01) and a high temporal resolution one (EPI02). Parameters for EPI01 were: FOV = 15 × 10.05 mm^2^, matrix size = 100 × 67, in-plane resolution = 0.15 × 0.15□mm^2^, slice number = 15, slice thickness = 0.4 mm, flip angle = 60°, TR/TE = 1500 ms /15 ms, repetitions = 256. EPI02 images were acquired with the following parameters: FOV = 15 × 12 mm^2^, matrix size 75 × 60, in-plane resolution = 0.2 × 0.2 mm^2^, slice number = 10, slice thickness = 0.4 mm, flip angle = 35°, TR/TE = 350 ms/15 ms, repetitions = 1100.

### 2.2 U-Net

The U-Net architecture is shown in Figure 1. It consists of one encoding and one decoding path with a skip connection. The skip connection, which connects the corresponding downsampling and upsampling stages, allows the model to integrate multi-scale information and better propagate the gradients for improved performance. Notably, the anatomical and functional images were usually acquired slice-by-slice with 2D sequence to span the whole brain in the fMRI study, resulting in an intra-slice spatial resolution that is significantly higher than inter-slice spatial resolution. In such case, the conventional approach is to take the individual 2D slice as input and predict the corresponding brain mask with 2D U-Net. However, the 2D model ignores the inter-slice information. To capture more spatial feature information while preserve intra-slice information, we took the multi-slice 2D images as an anisotropic 3D volume and used 3D U-Net model for training by only maxpooling and upsampling within the slice.

**Figure 1.**
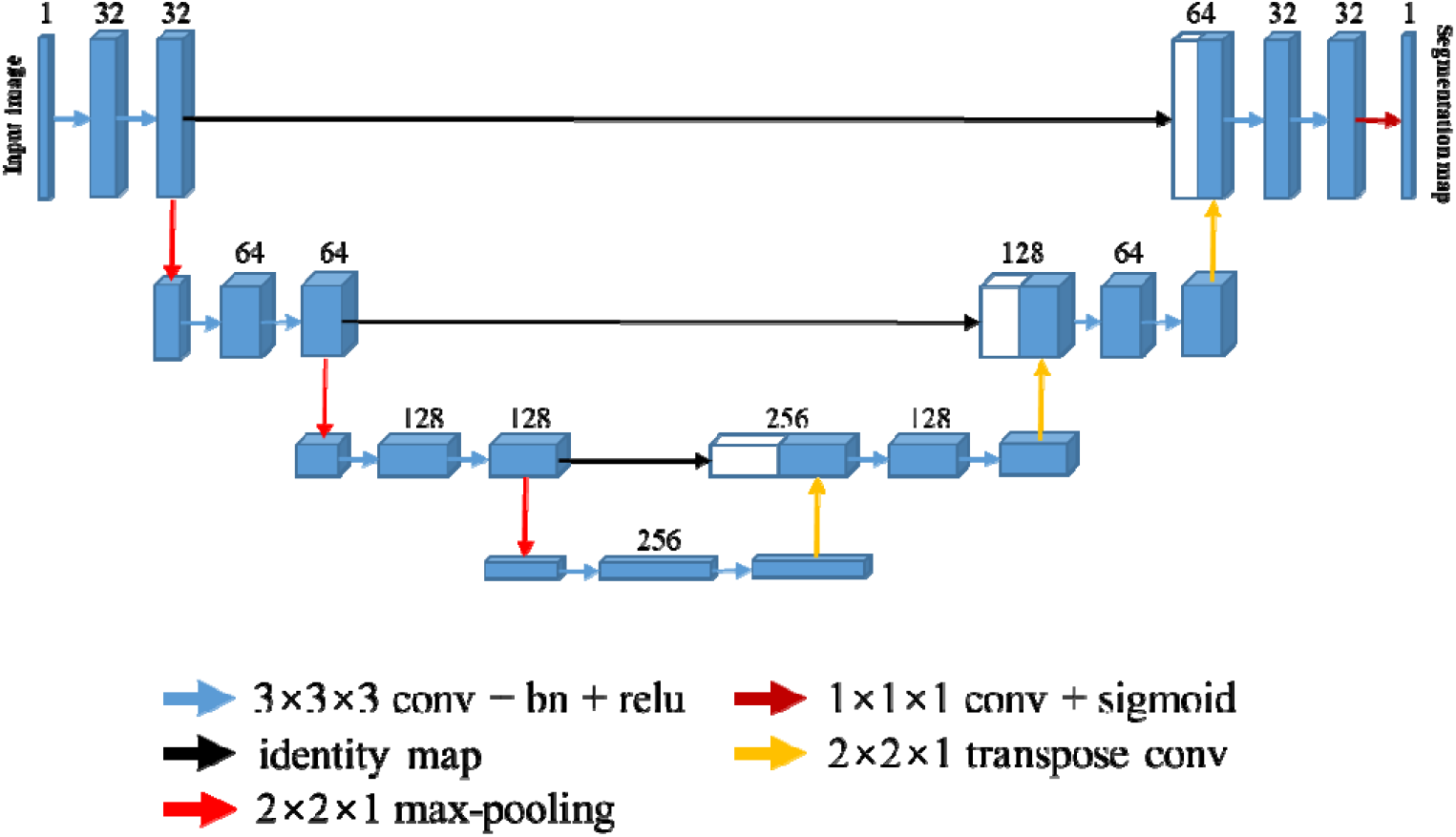
Architecture of the 3D U-Net for skull stripping. Each blue box indicates a multi-channel feature map, and the number of channels is denoted on top of the box. Every white box indicates the copied feature map. Color-coded arrows denote the different operations.

Every stage in both the encoding and decoding path is composed of two repeated 3 3 3 convolutions, each of which is followed by a rectified linear unit (ReLU), and a 2 2 slice max-pooling layer (in the encoding path) or a 2 × 2 slice deconvolutional layer (in the decoding path). The number of downsampling and upsampling is set to three, and the channel number of first stage is set to 32, being doubled at each downsampling step and then halved at each upsampling step. The output layer following the last decoding stage is a 1 1 1 single channel convolutional layer with a sigmoid activation function, which transforms the feature representation to one segmentation map.

Considering that the appearances of T2w images were significantly different from those of T2*w images, using the data from both modalities to train a network may degrade the segmentation performance. In this study, two models were trained separately with anatomical T2w images and functional T2*w images based on the above 3D U-Net architecture. As functional T2*w data include multiple repetitions in each section, only the first repetition was used in our study. A total of 74 mice were randomly selected from the first dataset for training, and the remaining 10 mice were used for testing. All of the second dataset were also used as test data to verify the effectiveness of our model across different data sources. In the training phase, 80% of training data from 74 mice were selected randomly to train the model, and the remaining 20% was used to validate the model to avoid over-fitting. Furthermore, data augmentation like flipping and rotating around three axes was also used to increase the diversity of the training dataset to improve the generalization of the models.

The 3D U-Net models was implemented using TensorFlow1.12.0 (Abadi et al., 2016) and trained on a Nvidia Titan X GPU (12GB). The convolution parameters were randomly initialized from a normal distribution with mean value of zero and a standard deviation of 0.001. Adam was used to optimize the training network (Kingma and Ba, 2014) with mini-batch size of two, and batch normalization (Ioffe and Szegedy, 2015) was adopted to accelerate the network training. The maximal number of training epochs was set to 100. The learning rate started from 10^−4^ and decayed by a factor of 0.99 every epoch. The loss function used to train the models was focal loss (Lin et al., 2020), which is designed to address class imbalance and can focus training on a sparse set of hard examples with large errors.

### 2.3 Comparison of methods

3D U-Net models were compared to two widely used methods for mouse brain extraction: RATS and SHERM. The parameters of both methods were carefully tuned to achieve the best performance for each dataset. For RATS (Oguz et al., 2014), the brain volume was set to 380 mm^3^ for both D1 and D2. In D1, the intensity threshold was set to 2.1 times of average intensity of each image for the T2w modality and 1.2 times for the T2*w modality. In D2, the threshold value was set to 1.3 times for the T2w and 0.6 times for the T2*w. For SHERM (Liu et al., 2020), the range of brain volume was set to 300–550 mm^3^. Except for the above parameters, all others were set to the default values for RATS and SHERM.

### 2.4 Data pre- and post-processing

All images were preprocessed before being fed into the models as follows. First, the N4bias field correction (Tustison et al., 2010) was applied to correct the intensity inhomogeneity for all images with the Python SimpleITK (Yaniv et al., 2018). Second, we resampled all data to a resolution of 0.0625 × 0.0625 mm^2^ for the anatomical T2w images and 0.25 × 0.25 mm^2^ for the functional T2*w images within the slice, leaving the inter-slice resolution unchanged. Then, all images were zero-padded to a size of 256 × 256 for the anatomical images and 64 × 64 for the functional images within slices. The padding was not performed in the inter-slice direction when the slice number was larger than 20, otherwise the slice number was padded to 20 with edge values of the slice. Subsequently, the histogram of each image was matched to a target histogram that was the average of all histograms of the training dataset. Finally, each anisotropic volume data was normalized in the range [0, 1].

After obtaining the binary mask from the network’s probability output, the only post-processing step was to identify the largest connected component and discard all the others (disconnected ones) for the final brain mask.

### 2.5 Evaluation methods

Two types of assessment methods were used to evaluate the effectiveness of the 3D U-Net models. The first type measured the overlap of the predicted segmentation mask (*M*_*predicted*_) generated by each skull-stripping algorithm and the manual segmentation mask (*M*_*manual*_). The quantify metrics include the Dice coefficient, the Jaccard index, and the Hausdorff distance.

The Dice coefficient is defined as twice the size of the intersection of the two masks divided by the sum of their sizes:

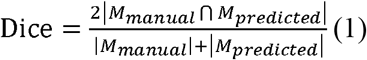

The Jaccard index is defined as the size of the intersection of the two masks divided by the size of their union:

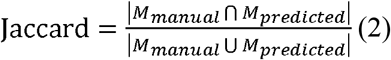

The Hausdorff distance (Huttenlocher et al., 1993) between two finite point sets is defined as:

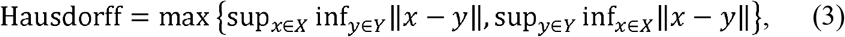

where *X* and *Y* denote the boundaries of the predicted segmentation mask and the manual segmentation mask, respectively. To exclude possible outliers, the Hausdorff distance is redefined as the 95th percentile distance instead of the maximum in our study.

The second assessment involves evaluating the impact of automatic skull stripping with 3D U-Net models on the final result of the fMRI analysis. The fMRI data were preprocessed with the common pipeline after skull stripping and then analyzed with seed-based analysis (Chan et al., 2017; Wang et al., 2019) and group independent component analysis (group ICA) (Mechling et al., 2014; Zerbi et al., 2015), which are two widely used methodologies in fMRI studies. Seed-based analysis requires creating a seed region first and then generates a seed-to-brain connectivity map by calculating the Pearson’s correlation coefficient (CC) between the BOLD time course of the seed and those of all other voxels in the brain. Group ICA is a data-driven method for the blind source separation of fMRI data, which is implemented in the group ICA of fMRI toolbox (GIFT) (Rachakonda et al., 2007). The Pearson’s correlation coefficient was calculated between the time courses of each pair of components extracted from the group ICA.

## 3 Results

Figure 2 illustrates the segmentation results of RATS, SHERM, and the 3D U-Net model on the T2w images from one mouse in D1 (figure 2a) and from two mice of auditory (**b**) and somatosensory (**c**) stimulation in D2. As shown in Figure 2a, all three methods successfully extracted most brain tissue. By comparison, the segmentation contours of the 3D U-Net model and SHERM were smoother and closer to the ground truth than those of RATS. However, the brain mask predicted by SHERM misaligned with the ground truth at the sharp-angled corner, while the 3D U-Net model still performed well in these locations. As shown in Figure 2b and 2c, the segmentation performance on T2w images of D2 was degraded for RATS and SHERM, compared to that of D1. The performance degradation might be attributed to T2w images of D2 having lower SNR and image contrast than those of D1. Specifically, the RATS produced a brain mask with a rough boundary, while the SHERM evidently overestimated the entire brain volume on D2. However, the 3D U-Net model still performed well on both D2 and D1.

**Figure 2.**
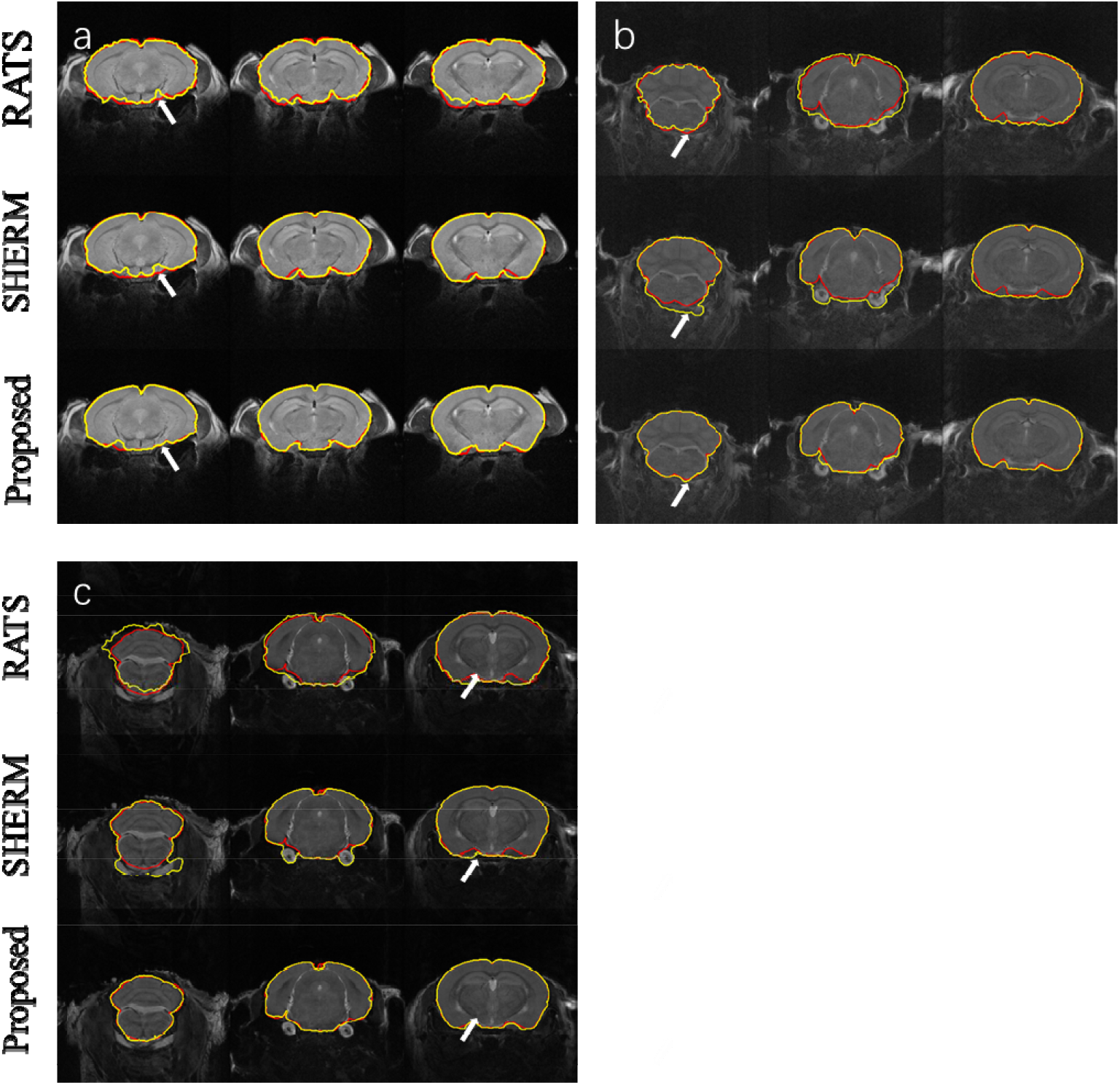
Example segmentation comparison for T2w images from one mouse in D1 (**a**) and from two mice with auditory (**b**) and somatosensory (**c**) stimulation in D2. Red lines show the contours of ground truth; yellow lines show automatically computed brain masks by RATS, SHERM, and the 3D U-Net model. White arrows point to the rough boundary, where the 3D U-Net model performed better than RATS and SHERM.

Figure 3 shows the segmentation results for three representative slices of RATS, SHERM, and the 3D U-Net model on T2*w images from one mouse in D1 (Figure 3a) and from four mice in D2 with high spatial resolution (EPI01) (Figure 3b_1, 3c_1) and high temporal resolution (EPI02) (Figure 3b_2, 3c_2). The SHERM overestimated the brain volume, especially for the D2. The RATS and 3D U-Net model performed better than SHERM in both D1 and D2. However, the brain boundaries predicted by the RATS were misaligned with the ground-truth boundaries in the areas with severe distortions or signal losses (white arrows), while the 3D U-Net model still had an accurate alignment on these locations.

**Figure 3.**
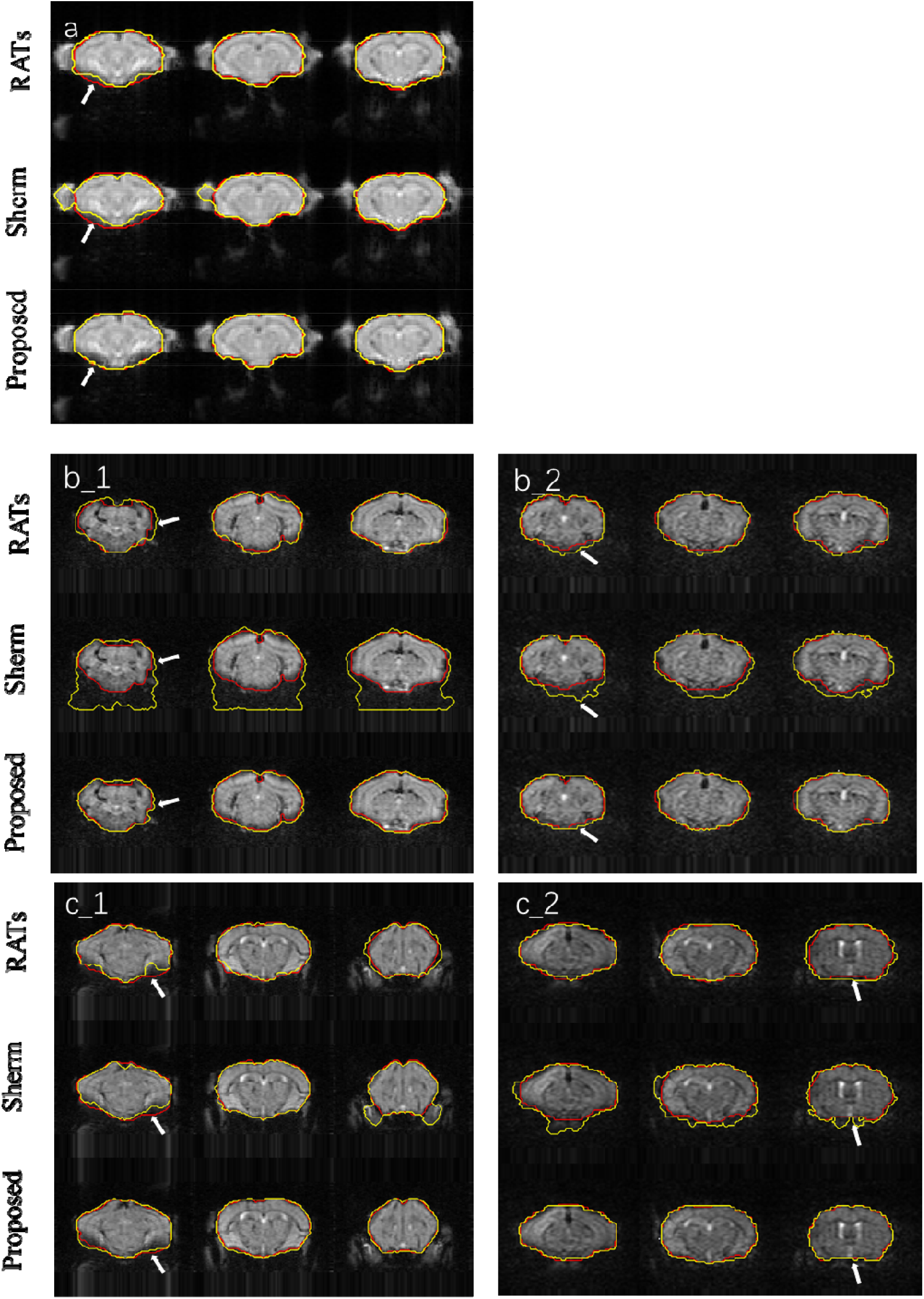
Example segmentations comparison for T2*w images from one mouse in D1 (**a**) and from two mice with auditory stimulation (**b**) and two mice with somatosensory stimulation (**c**) in D2; **b_1, c_1** represent EPI01, and **b_2, c_2** represent EPI02. Red lines show the contours of ground truth; yellow lines show automatically computed brain masks by RATS, SHERM, and the 3D U-Net model. White arrows point to the rough boundary, where 3D U-Net models performed better than RATS and SHERM.

Tables 1 and Table 2 show the quantitative assessment of RATS, SHERM, and 3D U-Net models for T2w images and T2*w images, respectively. The 3D U-Net models yielded highest mean values of the Dice and Jaccard index, and lowest mean values of the Hausdorff distance in both T2w and T2*w images. In addition, the standard deviations of all three metrics were lowest for the 3D U-Net models in most cases, except for the Hausdorff distance on T2w images of D2_PART2, and the Dice and Jaccard index on T2*w images of both D1 and D2_PART2_EPI01, which were slightly higher than those of RATS. The above quantitative results indicate that the 3D U-Net models exhibit a high accuracy and stability.

**Table 1.**
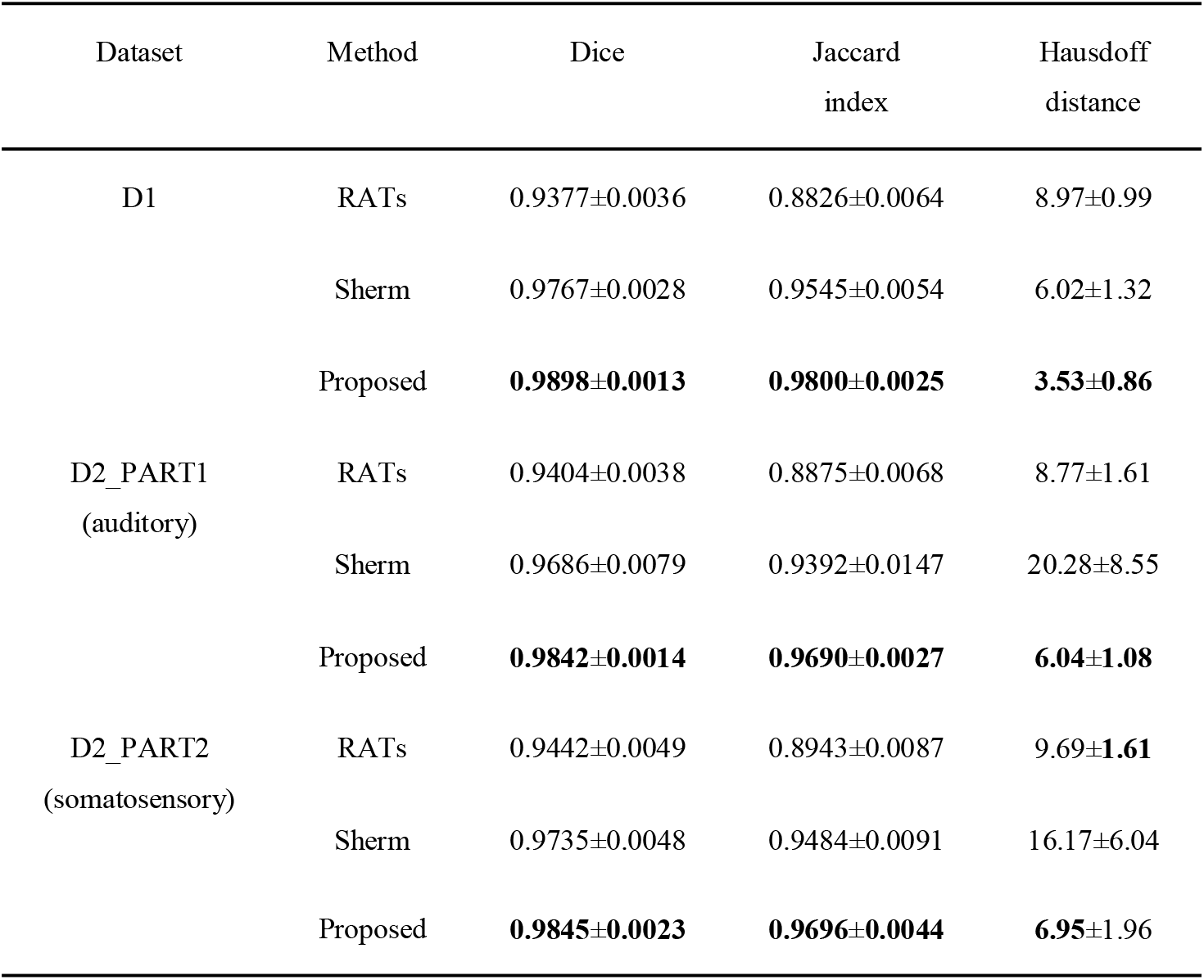
Mean and standard deviation of Dice, Jaccard index, and Hausdorff distance evaluating the RATS, SHERM, and 3D U-Net model for T2w images in different datasets. Bold values indicate the best results.

| Dataset | Method | Dice | Jaccard<br>index | Hausdorff<br>distance |
| --- | --- | --- | --- | --- |
| D1 | RATs | 0.9377±0.0036 | 0.8826±0.0064 | 8.97±0.99 |
|  | Sherm | 0.9767±0.0028 | 0.9545±0.0054 | 6.02±1.32 |
|  | Proposed | **0.9898±0.0013** | **0.9800±0.0025** | **3.53±0.86** |
| D2_PART1<br>(auditory) | RATs | 0.9404±0.0038 | 0.8875±0.0068 | 8.77±1.61 |
|  | Sherm | 0.9686±0.0079 | 0.9392±0.0147 | 20.28±8.55 |
|  | Proposed | **0.9842±0.0014** | **0.9690±0.0027** | **6.04±1.08** |
| D2_PART2<br>(somatosensory) | RATs | 0.9442±0.0049 | 0.8943±0.0087 | 9.69±**1.61** |
|  | Sherm | 0.9735±0.0048 | 0.9484±0.0091 | 16.17±6.04 |
|  | Proposed | **0.9845±0.0023** | **0.9696±0.0044** | **6.95±1.96** |

**Table 2.**
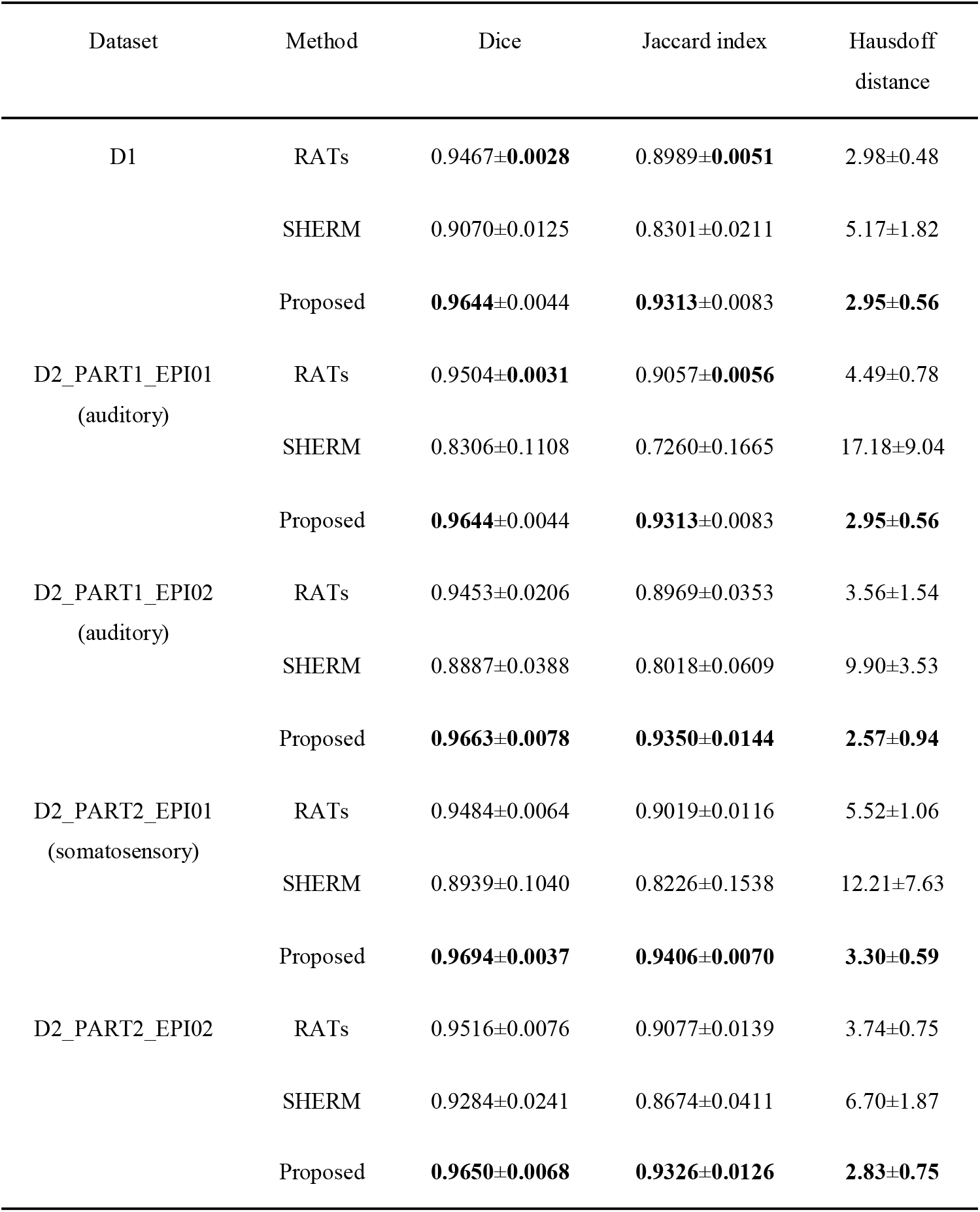
Mean and standard deviation of Dice, Jaccard index, and Hausdorff distance evaluating the RATS, SHERM, and 3D U-Net model for T2*w images in different datasets. Bold values indicate the best results.

| Dataset | Method | Dice | Jaccard index | Hausdorff distance |
| --- | --- | --- | --- | --- |
| D1 | RATs | 0.9467±**0.0028** | 0.8989±**0.0051** | 2.98±0.48 |
|  | SHERM | 0.9070±0.0125 | 0.8301±0.0211 | 5.17±1.82 |
|  | Proposed | **0.9644**±0.0044 | **0.9313**±0.0083 | **2.95±0.56** |
| D2_PART1_EPI01<br>(auditory) | RATs | 0.9504±**0.0031** | 0.9057±**0.0056** | 4.49±0.78 |
|  | SHERM | 0.8306±0.1108 | 0.7260±0.1665 | 17.18±9.04 |
|  | Proposed | **0.9644**±0.0044 | **0.9313**±0.0083 | **2.95±0.56** |
| D2_PART1_EPI02<br>(auditory) | RATs | 0.9453±0.0206 | 0.8969±0.0353 | 3.56±1.54 |
|  | SHERM | 0.8887±0.0388 | 0.8018±0.0609 | 9.90±3.53 |
|  | Proposed | **0.9663**±**0.0078** | **0.9350**±**0.0144** | **2.57±0.94** |
| D2_PART2_EPI01<br>(somatosensory) | RATs | 0.9484±0.0064 | 0.9019±0.0116 | 5.52±1.06 |
|  | SHERM | 0.8939±0.1040 | 0.8226±0.1538 | 12.21±7.63 |
|  | Proposed | **0.9694**±**0.0037** | **0.9406**±**0.0070** | **3.30±0.59** |
| D2_PART2_EPI02 | RATs | 0.9516±0.0076 | 0.9077±0.0139 | 3.74±0.75 |
|  | SHERM | 0.9284±0.0241 | 0.8674±0.0411 | 6.70±1.87 |
|  | Proposed | **0.9650**±**0.0068** | **0.9326**±**0.0126** | **2.83±0.75** |

Figure 4 compares the results of seed-based analysis from one mouse in the test data of D1 with automatic skull stripping by the 3D U-Net models and manual brain extraction. The seeds (2 □× □2 voxels) were positioned in the dorsal striatum (dStr), somatosensory barrel field cortex (S1BF), secondary somatosensory cortex (S2), and ventral striatum (vStr). The CC maps with our model were similar to those with manual brain extraction. We also plotted scatter plots, where the horizontal axis represents the values of CC maps with predicted masks, and the vertical axis represents the values of CC maps with the manual mask. Each point represents a pair of the two values at the same pixel location. All points were concentrated on the diagonal with R^2^ = 1. The above results indicate that the CC maps with automatic skull stripping by 3D U-Net models were identical to those with manual brain extraction.

**Figure 4.**
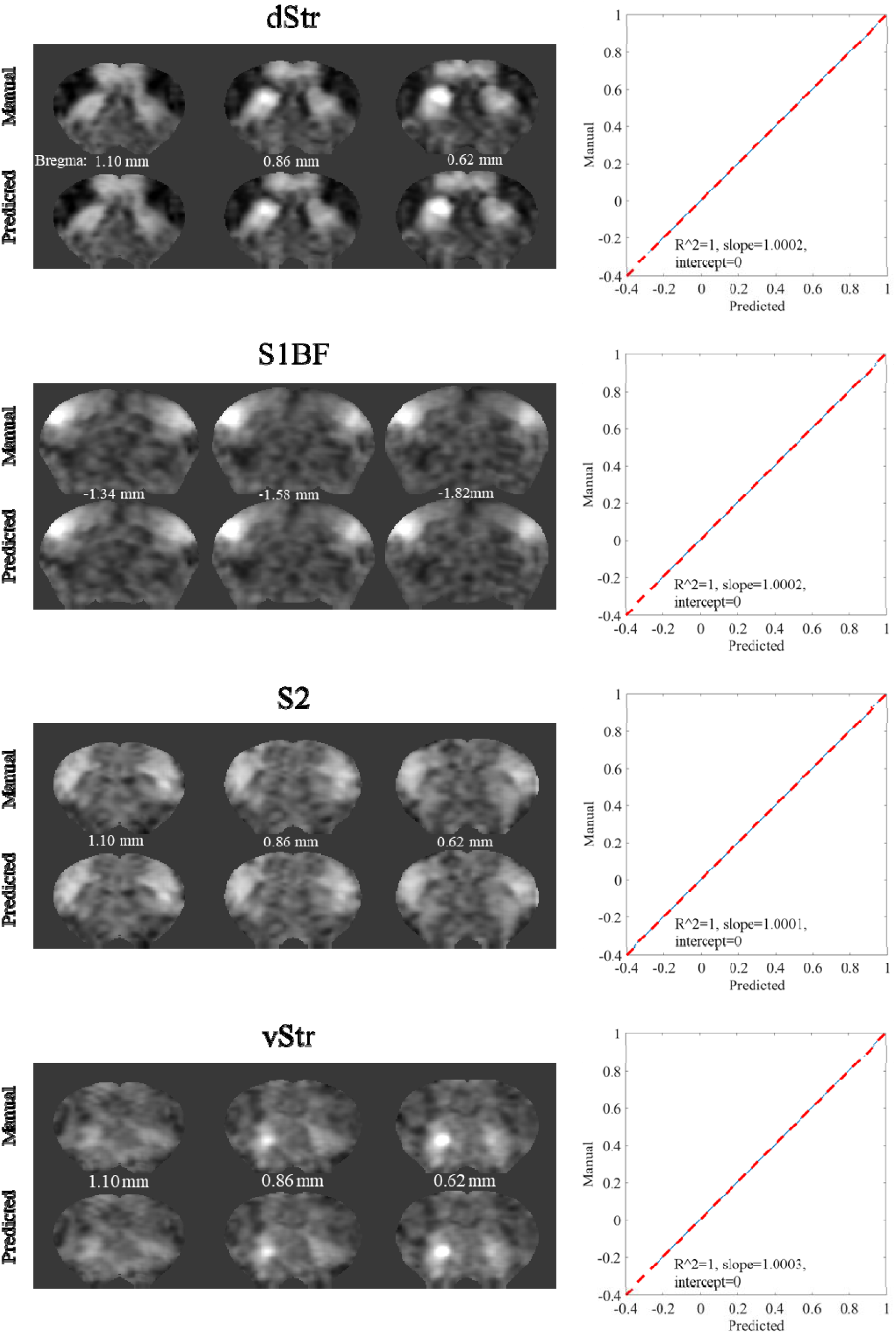
Exemplary results of seed-based analysis for one mouse in the test data of D1. The selected seed regions were S1BF, S2, dStr, and vStr. The left column illustrates the CC maps with manual brain extraction and automatic skull stripping by the 3D U-Net models for each seed region. The corresponding right column illustrates the scatters, where each point represents a pair of the two values from different CC maps with manual brain extraction and automatic skull stripping by the 3D U-Net models, at the same pixel location.

Figure 5 compared the four components of group ICA analysis with automatic skull stripping by the 3D U-Net models and manual brain extraction. The presented four components in Figure 5 correspond to the same regions used in the seed-based analysis (S1BF, S2, dStr, vStr). The component maps with 3D U-Net models for skull stripping were visually similar to those with manual brain extraction. The points in the scatter plots were concentrated on the diagonal with R^2^ = 1. These results indicate that the component maps of group ICA with automatic skull stripping by 3D U-Net models are highly consistent with those with manual brain extraction. Group ICA analysis with 3D U-Net models generated the same 32 independent components as those with manual brain extraction. The regions corresponding to each component are shown in Supplementary Table S1.

**Figure 5.**
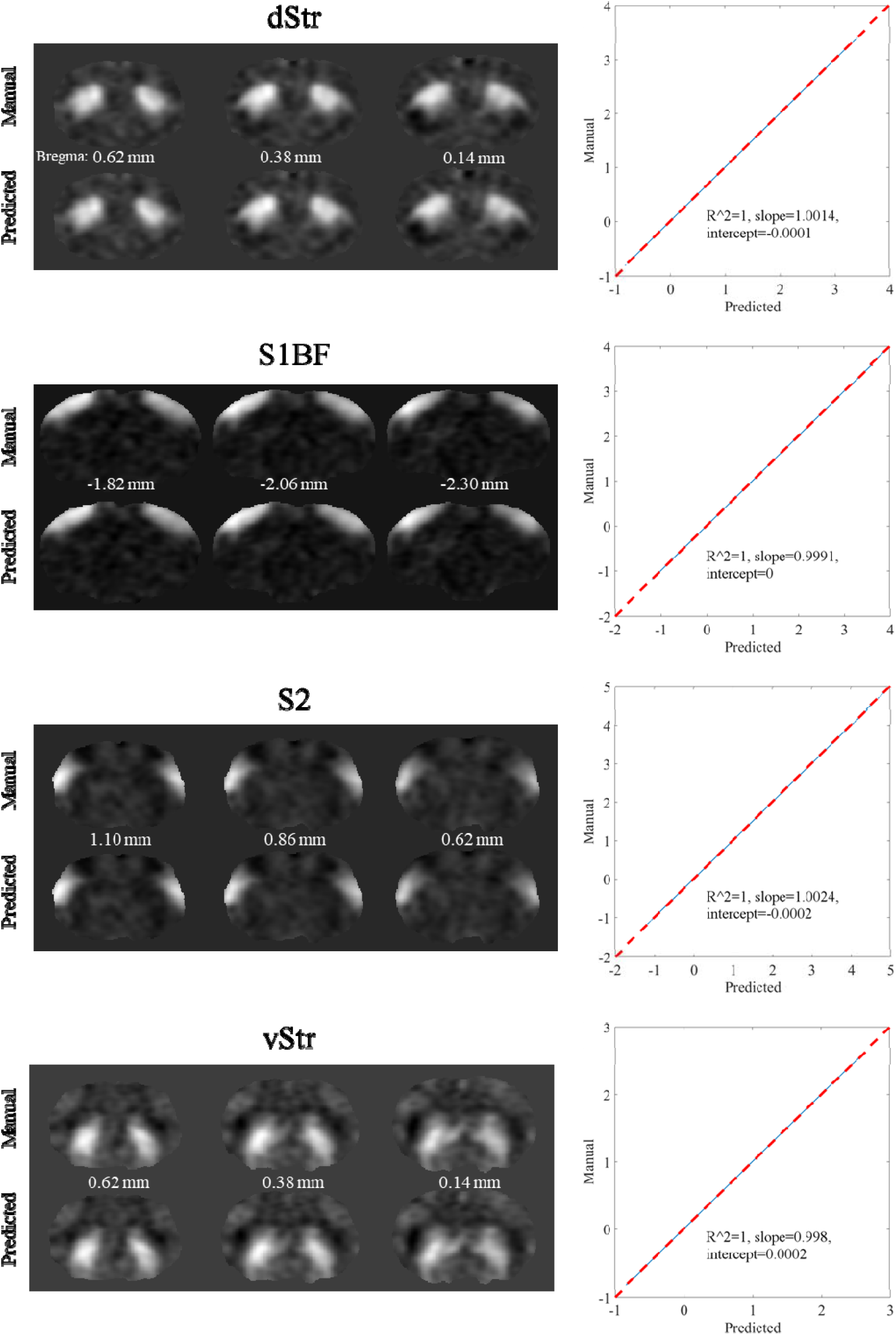
Example results of ICA analysis for test data of D1. Four selected components extracted by group ICA matching the four seed regions in seed-based analysis are shown in each row. The left column illustrates the component maps with manual brain extraction and automatic skull stripping by 3D U-Net models. The right column illustrates the scatters, where each point represents a pair of the two values from different component maps with manual brain extraction and automatic skull stripping by the 3D U-Net models, respectively, at the same pixel location.

Figure 6 shows the functional network connectivity (FNC) correlations for one mouse and the average FNC correlation across 10 mice between each pair of 32 regions extracted from group ICA analysis. Each correlation value in FNC matrixes with automatic skull stripping by 3D U-Net models was identical to that obtained with manual brain extraction. The points in the scatter plots are concentrated on the diagonal with R^2^ = 1, which indicates that the use of the 3D U-Net models for skull stripping in the fMRI analysis pipeline does not affect the final results of group ICA based FC analysis. The FNC correlations of the other nine mice are shown in Supplementary Figure S1.

**Figure 6.**
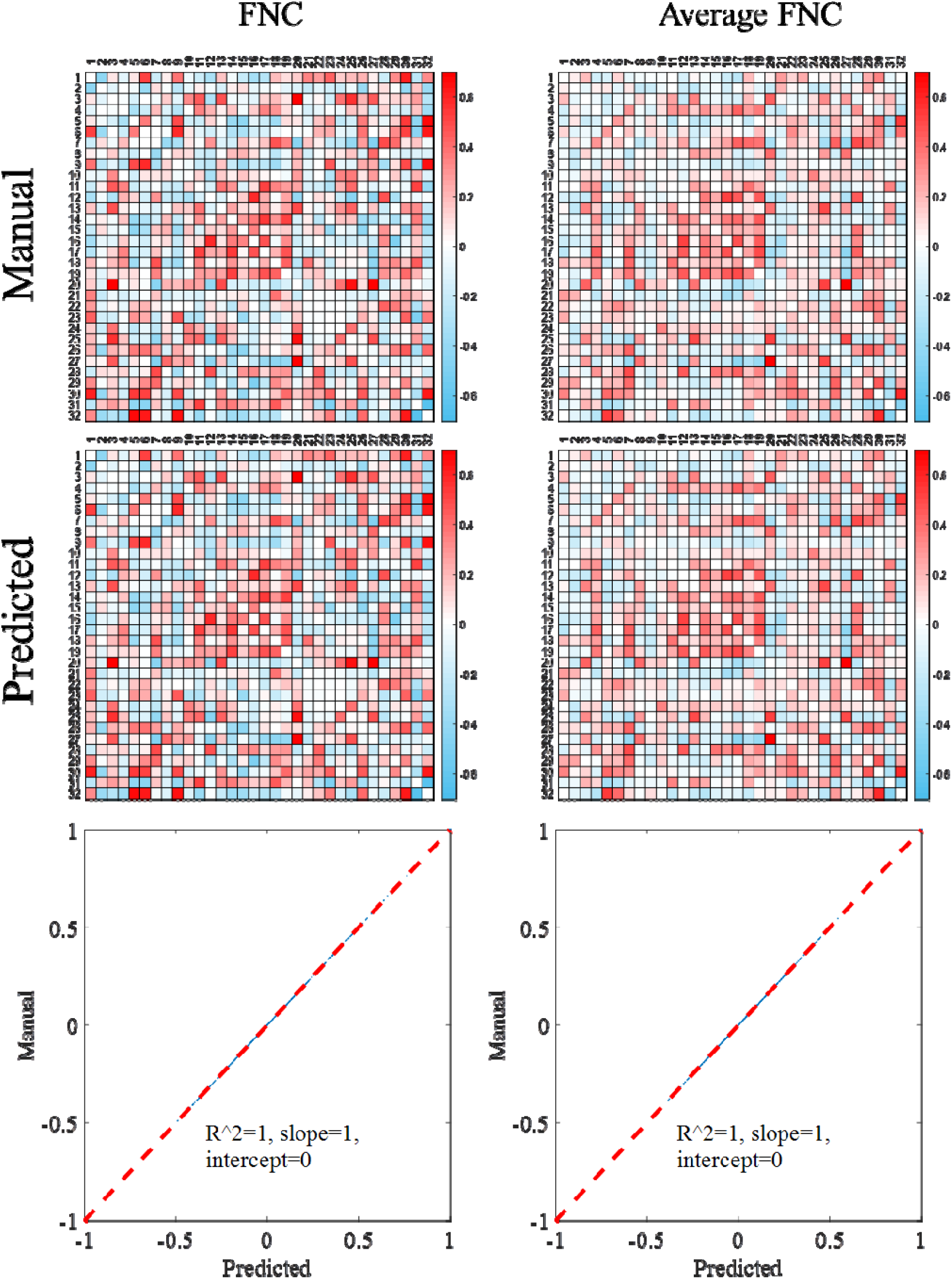
Functional connectivity between the independent components extracted from group ICA analysis for the test data of D1. The left column illustrates the FNC matrixes from one mouse, and the right column illustrates the average FNC matrixes across 10 mice. Scatter plots are shown in the bottom row, where each point represents the difference between two values from two FNC matrixes with manual brain extraction and automatic skull stripping by the 3D U-Net models, at the same pixel location.

## 4 Discussion

To the best of our knowledge, this is the first study investigating the feasibility of 3D U-Net for mouse skull stripping from brain functional (T2*w) images and the impact of automatic skull stripping on the final fMRI analysis. Results indicate that the 3D U-Net model performed well on both anatomical (T2w) and functional (T2*w) images and achieved state-of-the-art performance in mouse brain extraction. The fMRI results with automatic brain extraction using 3D U-Net models are nearly identical with those with manual brain extraction. Thus, the manual brain extraction in the fMRI pre-processing pipeline can be replaced by the proposed automatic skull-stripping method.

The 3D U-Net models were tested on not only interior but also exterior datasets. Notably, the exterior datasets were acquired with different acquisition parameters on a scanner with different field strength from another MRI center. The results show that 3D U-Net models had high segmentation accuracy that is comparable between interior and exterior datasets. This demonstrates that the developed method has high reliability and excellent generalization ability. The 3D U-Net models also outperformed two widely used brain extractions for rodent (SHERM and RATS). SHERM has the second best performance in T2w images (Figure 2), while it has the worst performance in T2*w images (Figure 3). The reason for this is that the poor quality of T2*w images makes it difficult to match the shape of the brain template. Although RATS has stable performance across different datasets and modalities, its segmentation accuracy (Dice < 0.945 in T2w images and Dice < 0.952 in T2*w images) is consistently lower than in our method (Dice > 0.984 in T2w images and Dice > 0.964 in T2*w images).

There are several related reports on using U-Net for mouse skull stripping (De Feo et al., 2021; Hsu et al., 2020; Thai et al., 2019). Compared with the model adopted by Hsu et al. (2020), the segmentation accuracy of our models is relatively higher in both T2w and T2*w images. The first reason is that we used 3D U-Net for mouse skull stripping, while Hsu et al. used 2D U-Net. The second reason is that we trained the U-Net models separately for T2- and T2*w images. The performance of our 3D U-Net model is comparable to that of the 3D model adopted by De Feo et al. (2021) on T2w anatomical images. We also applied the 3D U-Net for brain extraction from T2*w functional images. The multi-task U-Net developed by De Feo et al. can hardly be applied to functional images, because it is difficult to delineate different brain regions in functional images due to their low spatial resolution, contrast, signal-to-noise ratio, and severe distortion.

It is essential to guarantee that automatic skull-stripping method does not alter fMRI analysis results. Thus, we not only evaluated the segmentation accuracy, but also investigated the effect of automatic segmentation on fMRI analysis results. The fMRI analysis results with automatic skull stripping by 3D U-Net models are identical to those with manual skull stripping. This finding demonstrates that the 3D U-Net based method can replace manual skull stripping and facilitate the establishment of the automated fMRI analysis pipeline for the mouse model.

With respect to the computational cost, the 3D U-Net based method proves to be time efficient. The computation time of the 3D U-Net method was approximately 3 s for a T2w volume data with a size of 256 × 256 × 20, and 0.5 s for T2*w volume data with a size of 64 × 64 × 20. In comparison, the computation time of SHERM is 780 s and 3 s for T2w and T2*w images, respectively; the computation time of RATS is 10 s for T2w images and 3 s for T2*w images. All test procedures were run on a server with a Linux 4.15.0 system, an Intel(R) Xeon(R) E5-2667 8-core CPU, and 256 GB RAM.

There are two limitations in our current work. First, the segmentation accuracy of the developed 3D U-Net model on functional images is still relatively lower than that on anatomical images, because of the poor image quality of the functional images. Utilizing the cross-modality information between anatomical and functional images may further improve the accuracy of skull stripping on functional images. Second, the developed 3D U-Net model was only trained and validated on adult C57BL/6 mice, and cannot be directly applied to brain MR images from different mouse types and ages. To address this problem, the model need to be retrained by including more manually labeled data from mice with varying types and ages. Labeling data is time-consuming and labor-intensive, and another potential approach to reduce the amount of labeled data is to utilize transfer learning (Long et al., 2015; Yu et al., 2019; Zhu et al., 2021).

## 5 Conclusion

We investigated an automatic skull-stripping method based on 3D U-Net for mouse fMRI analysis. The 3D U-Net based method achieves state-of-the-art performance on both T2w and T2*w images in terms of the segmentation accuracy. Identical results between mouse fMRI analysis using manual and automatic skull stripping demonstrates that the 3D U-Net model has a great potential to replace manual labeling in the mouse fMRI analysis pipeline. Hence, skull stripping by the 3D U-Net model will facilitate the establishment of an automatic pipeline of mouse fMRI data processing.

## Acknowledgments

This work receives support from the National Natural Science Foundation of China (61971214, 81871349); the Natural Science Foundation of Guangdong Province (2019A1515011513); the Guangdong-Hong Kong-Macao Greater Bay Area Center for Brain Science and Brain-Inspired Intelligence Fund (2019022);the Technology R&D Program of Guangdong (2017B090912006); the National Key R&D Program of China (2019YFC0118702).

