## Supplementary material for "Automated skull stripping in mouse fMRI analysis using 3D U-Net": Figure S1

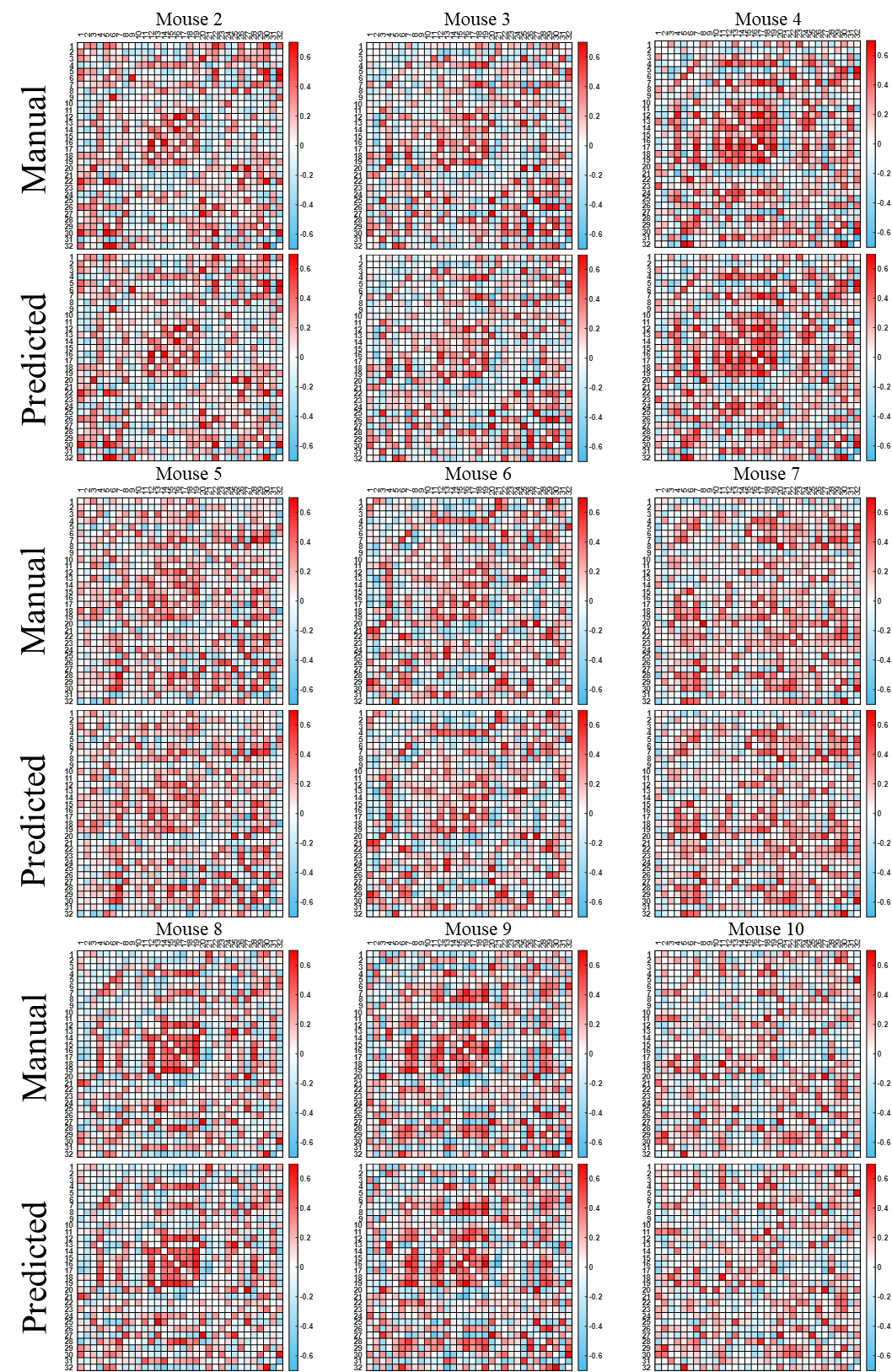

Figure S1 - Functional network connectivity (FNC) matrixes of the remaining nine mice with automatic skull stripping by 3D U-Net models and using manual brain extraction.
