## Supplementary material for "Automated skull stripping in mouse fMRI analysis using 3D U-Net": Table S1

Table S1 - A total of 32 components extracted by the group ICA analysis and corresponding brain regions.

| Component | Region | Component | Region |
| --- | --- | --- | --- |
| 1 | anterior lobe cerebellum; subiculum; presubiculum; parasubiculum | 17 | primary somatosensory cortex, barrel field; primary somatosensory cortex; primary visual cortex; secondary visual cortex, lateral part; primary somatosensory cortex; secondary auditory cortex, dorsal part |
| 2 | lateral ventricle | 18 | ventral orbital cortex; lateral orbital cortex; medial orbital cortex |
| 3 | glomerular layer of the olfactory bulb; dysgranular insular cortex | 19 | secondary motor cortex |
| 4 | dorsal striatum | 20 | primary motor cortex |
| 5 | thalamus | 21 | prelimbic cortex; infralimbic cortex |
| 6 | 4 & 5 cerebellar lobules | 22 | external cortex of the inferior colliculus; simple lobule |
| 7 | cingulate cortex; secondary motor cortex | 23 | field CA1 of hippocampus; field CA2 of hippocampus; field CA3 of hippocampus; dentate gyrus; subiculum; presubiculum |
| 8 | primary somatosensory cortex, forelimb region; primary somatosensory cortex, hindlimb region | 24 | primary visual cortex |
| 9 | dorsal nucleus of the lateral lemniscusk; intermediate nucleus of the lateral lemniscus | 25 | medial striatum |
| 10 | primary visual cortex | 26 | retrosplenial granular cortex |
| 11 | ventral striatum | 27 | primary somatosensory cortex, barrel field |
| 12 | primary somatosensory cortex, forelimb region; primary somatosensory cortex, hindlimb region | 28 | primary somatosensory cortex, trunk region |
| 13 | secondary somatosensory cortex; secondary auditory cortex, ventral part; temporal association cortex; ectorhinal cortex; perirhinal cortex | 29 | Cingulate cortex |
| 14 | primary somatosensory cortex, barrel field | 30 | superior colliculus |
| 15 | secondary visual cortex, lateral part; secondary auditory cortex, dorsal part; primary auditory cortex; secondary auditory cortex, ventral part | 31 | basolateral amygdaloid nucleus, anterior part; basomedial amygdaloid nucleus, anterior part; medial amygdaloid nucleusmanterior part |
| 16 | primary somatosensory cortex, forelimb region; primary somatosensory cortex, hindlimb region | 32 | superior colliculus |
